# Exploring Transcriptional Regulation of Soybean Tissue Development with Machine Learning Method

**DOI:** 10.1101/2024.08.12.607582

**Authors:** Yong Yang

## Abstract

Soybean is one of the most important crops that is widely demanded by people in daily lives. Measuring the transcriptome of a tissue or condition is a powerful way to detect changes in genetic adaptation. However, it remains difficult to identify the key genes in transcriptional regulation most likely to explain specific traits. Here, we outline a machine learning method that utilizes publicly available soybean RNA-seq data by uncovering conserved expression patterns of genes controlled by transcription factor (TF) / transcription regulator (TR) genes in soybean tissues across time and space under various conditions. In addition to its function in gene expression homeostasis, we can also identify important TF/TR genes related to soybean leaf, stem and root tissue development. Combining with co-expression modules highly expression in the tissue, we also highlight the impact of candidate TF/TR genes in the module in different tissues that may shape the dynamics of soybean development. Together, our results revealed the importance of transcriptional regulatory module analysis in unraveling key traits in the soybean development, in particular those TFs/TRs and their target genes.

## Introduction

Soybeans are one of the world’s most important and highly nutritious crops, providing oil and protein for human beings and livestock. As a paleopolyploid plant, soybean had undergone two rounds of large-scale genome duplication, thus most of soybean genes were paralogous genes with multiple copies ^1^. Therefore, mining the key genes related to agronomic traits is of great significance for this genetically complex crop. With the development of high-throughput sequencing technologies, exploring the mRNA expression has becoming useful tools to understand underlying mechanisms in soybean nutritional components, abiotic signaling and organ development studies ^2-4^. Thus, changes in gene expression are thought to play an important role in the complex traits exhibited by plants adapting to the environment and conditions ^5^.

Transcription factors (TFs) played central roles to different cellular functions, and the combination of multiple TFs on DNA elements drove the expression of target genes and controlled the gene regulatory processes ^6^. TFs affected gene expression not just by acting on individual downstream genes, but by regulating the co-expression of a group of genes ^7^. Groups of genes expressed in cells were often co-regulated, in that they are regulated by the same regulatory processes ^8^. A common set of TFs can bind the promoters or enhancers of the downstream regulatory genes to ensure the coordinated process. Traditionally, only differentially expressed genes had been focused on and complemented by the cluster and enrichment analysis, genes which were not differentially expressed in the same interaction module were often ignored in transcriptional studies ^9^. Thus, a systems-level analysis to explore these modules remains to be performed.

Co-expression networks make it possible to discover important modules of genes which control the same biological process ^10^. These model clusters are annotated as functional or cellular specific processes, and also can be used to annotate external gene sets ^11, 12^. Thus, researchers had identified hub modules by conducting a gene co-expression network analysis by using transcriptome from different tissues and development stages in soybean^13, 14^. Soybean transcriptome data had multiplied in the past few years, and the increasing transcript sample size could enhance the statistical power to obtain more precise and robust molecular marker estimates and to reduce noise effects and individual study bias ^15^. Publication of new linear model provides important tool for mining key gene regulatory modules in model species ^16, 17^. Moreover, recent progress in machine learning models had showed the success in the field of biology and could facilitate the discovery and analysis of biological networks, which offered potential for non-model species ^18, 19^.

Here we constructed a neural network to accurately reconstitute transcriptome profiles based on the expression of TF/transcription regulators (TR) genes in a non-model plant, soybean. The expression of all other genes could be predicted by the combinatorial control induced by the TFs/TRs. Meanwhile, it can also be applied to other tissues. Then, xgboost model was trained to classify three different tissues based the TF/TR genes. This method ranked important TF/TR genes that is likely more able to prioritize functionally significant changes. Using the gene co-expression network modules in each tissue, we identified the specificity of the gene modules associated with tissue-related. We highlighted divergence in the expression of candidate TF/TR genes that may have played an important role in soybean tissue development and function.

## Method

### Data collection and processing

All the RNA-Seq expression data of soybean were collected from the Soybean Expression Atlas v2 database ^15^. We selected expression data from tissues with a sample size greater than 50 in soybeans for analysis. The transcripts per million (TMP) values were used for each gene. We first removed genes with low expression levels that had a TMP ≥ 1 in less than 500 samples. Then samples containing more than 20,000 genes with zero expression levels are removed. The expression values of 42,492 genes in 5,293 RNA-Seq sample were kept and merged into a large expression matrix. We separated the data into two sets, one of the genes annotated as soybean TF/TR gene set while were extracted according to the SoybeanGDB database ^20^, and one containing the rest of the genes. The rest genes were assumed to be regulated downstream of those TFs/TRs. Two datasets were log transformed via log^2^ (TPM +1) for downstream analysis.

### Model design

We next aimed to predict the expression levels of the non-TF/TR genes using the TF/TR expression levels. To this end, we designed the models to be feed-forward, fully connected neural networks. We built the models to have one input node for each TF/TR, totaling 3,704 input nodes and one output node for each target gene in the data, totaling 38,788 output nodes. Moreover, we used the Adam algorithm to minimize the Normalized Root Mean Squared Error (NRMSE), with a learning rate of 0.001, β1 = 0.9, β2 = 0.999, and decay of 0.01. Dropout layer was used to avoid overfitting ^21^. The whole model was optimized by weighting binary cross-entropy loss in different tasks. The model was trained in Keras using Python 3, and the performance was calculated using spearman correlation coefficients (r) between the true and the predicted expression levels of the non-TF/TR genes.

### Hyperparameter optimization and model selection

In order to obtain models with better performance and generalization capability, we performed hyperparameter optimization. We used the Tree of Parzen estimators (TPE) strategy to conduct hyperparameter searches using Hyperopt package ^22^. We trained a basic model by selecting a set of hyperparameters including hidden layers, hidden nodes, drop probability, batch size and activation functions. The performance of the predictor was evaluated using a 10-fold cross validation. The samples were randomly split into 10 equal folds. All samples except those in one-fold were used to train the model, while the remainder were used to valid the model. The average model performance with the accuracy in the validation set was calculated. It should be pointed out that only the combination of hyperparameters with the highest accuracy in the validation set was retained.

### Test data processing

Raw SRA data from four different soybean tissues: root (n=12), leaf (n=6), flower (n=3) and pod (n=3) under different treatments or conditions were download from three recently public researches ^23-25^. The raw reads were trimmed with fastp ^26^ to obtain clean data, and the resultant reads were mapped to soybean genome (Wm82.a4.v1, STAR version=2.7.10b ^27^). The genome annotation file was download from PLAZA Dicots 5.0 platform ^28^. Then the TPM values of all the genes were analyzed with the same pipeline as described above and used to test the performance of our predicting model. Moreover, transcriptome data from endosperm (n=10), epicotyl (n=6), petiole (n=6) and radicle (n=7) tissues were used to validate the model’s generalization ability ^15^.

### Construction classification machine learning model

We used a tree model with gradient boosting, XGBoost ^29^, to train and test the model to classify samples of root, shoot and leaf with a sample size of more than 1000. We used 10-fold internal cross-validation to select the optimized hyperparameters. We tuned “colsample_bytree”, “max_depth”, “min_child_weight”, “subsamples”, and “eta” in the XGBoost classification model. Subsequently, we made predictions on each of the left-out set, assessed the model accuracy, precision, recall, and F1 score by calculating the accuracy between the predicted and actual class.

### Gene modules identification

The gene module networks were inferred using the R package WGCNA ^30^. The expression matrix in the 12 tissues were used for co-expression network construction. We first assessed the scale-free topology fit to choose the most suitable β power (R^2^=0.8). The pairwise correlations between genes were first calculated, and then the correlation matrix was transformed into a an adjacent matrix which was used to construct the topological overlap matrix (TOM) ^31^. Then each TOM was used as input for hierarchical clustering analysis based on the dissimilarity (1-TOM), and gene modules was detected by using a dynamic tree cutting algorithm (deep split = 4 and minModuleSize = 30) ^32^. Finally, we calculated pairwise similarity between modules and merged the ones that had PCC ≥ 0.8. Correlations between these co-expression modules and soybean tissue characteristics were also identified. To further investigate these modules, the genes in these modules were performed for Gene Ontology enrichment using AgriGO with a q-value < 0.05 as significantly ^33^. Then we used an efficient algorithm for regulatory network inference GRNBoost2 to infer the TFs/TRs regulating the gene features ^34^.

## Results

In a typical RNA-seq pipeline, DEGs are identified by comparing the RNA-seq read counts between different conditions using a p-value and FDR for differential expression. Various enrichment knowledge then be used to identify interesting pathways and candidate genes. However, these previous methods do not consider co-regulatory effect in gene expression levels. To systematically apply this concept, we first predicted expression levels of non-TF/TR genes by the TF/TR genes (**Figure 1**). We use data from a large publicly available RNA sequencing dataset of a non-model species, soybean, to study its regulatory network. In addition, the expression levels of different TFs/TRs reflect the developmental processes and functions of different tissues and may better reflect the potential of gene expression changes to alter phenotypes. The ranked list is then used in conjunction with the co-expression network to determine the regulatory module for TFs/TRs and target genes.

**Figure 1.**
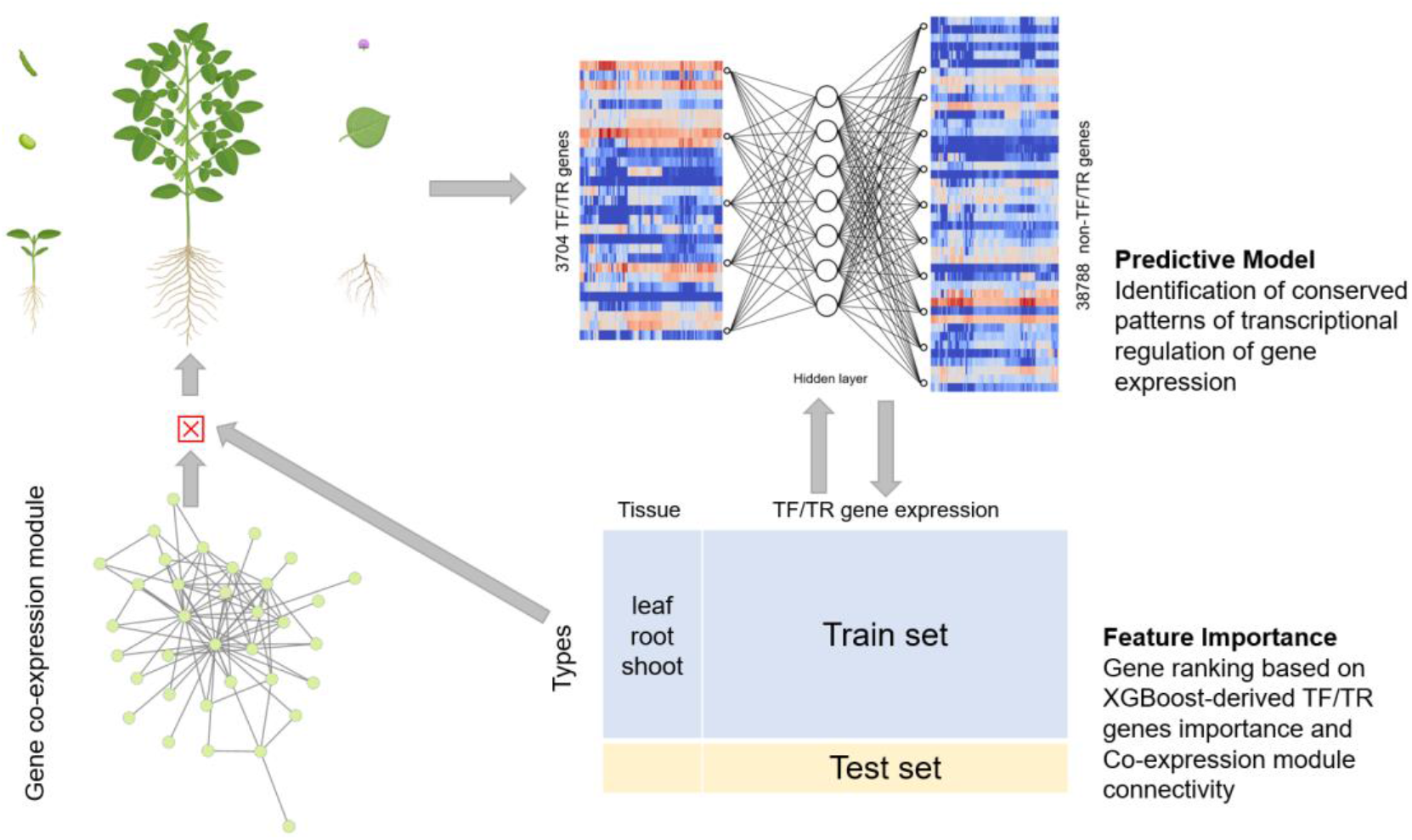
Outline of the methodology. Step 1 Soybean gene expression neural networks predictor: the network was made to predict the expression of non-TF/TR genes based on TF/TR genes. Step 2 feature importance: We ranked the TF/TR genes based on the XGBoost-derived feature importance score for different tissues. This biologically principled approach to reduce the TF features ultimately improved the predictor performance. Step 3 TF regulatory network: We extracted the tissue co-expression modules and combined with important TFs to construct regulatory network based on the TF connectivity by using GRNBoost2.

### Predicting soybean transcriptomes with TFs/TRs

We collected a total of 5,293 samples from 12 different tissues or growth stages. The samples clustered well by tissues in the t-SNE representation using TF/TR genes (**Figure S1**), and the heatmap of TF/TR gene expression clustering in different tissues also shows that TFs/TRs can also be well represented in different tissue clusters of the soybean using the average expression of each TF/TR (**Figure S2**). The predictive model of soybean gene expression was designed based on the premise that TFs integrated developmental and environmental cues to fine-tune the expression of their target genes and ultimately precise prediction of the expression level of the transcriptome. Thus, we applied the expression level of 3704 TF and TR genes including 90 gene families such as MYB, WRKY, and NAC gene family as the input data (**Figure S3**), fed to a fully connected neural network, and took the expression of 38788 non-TF/TR genes as the output feature. We used 1–4 intermediate layers with a depth of 50–1024 hidden nodes in each layer. Specifically, we conducted 10-fold cross-validation to evaluate the predictor’s performance and stability, in which the model was trained on 90% of the samples and tested on the remaining 10%. The rationale is to identify the most compact architecture of TFs/TRs to predict most of the target gene expression with sufficient accuracy. Among the, the one-layer model has the best performance, and the average correlation coefficient and NRMSE of each test samples were 0.9326 and 0.2062, while the two-layer were 0.9289 and 0.2155, three-layer were 0.9072 and 0.3148, and four-layer 4 were 0.8467 and 0.3351 (**Figure 2A, 2B**). We can observe that the accuracy decreases when the depth of the network increases. Based on Occam’s Razor’s Law, we choose the first layer of the model for subsequent analysis. The result showed that the model can achieved high accuracy in different tissues (**Figure 2C**).

**Figure 2.**
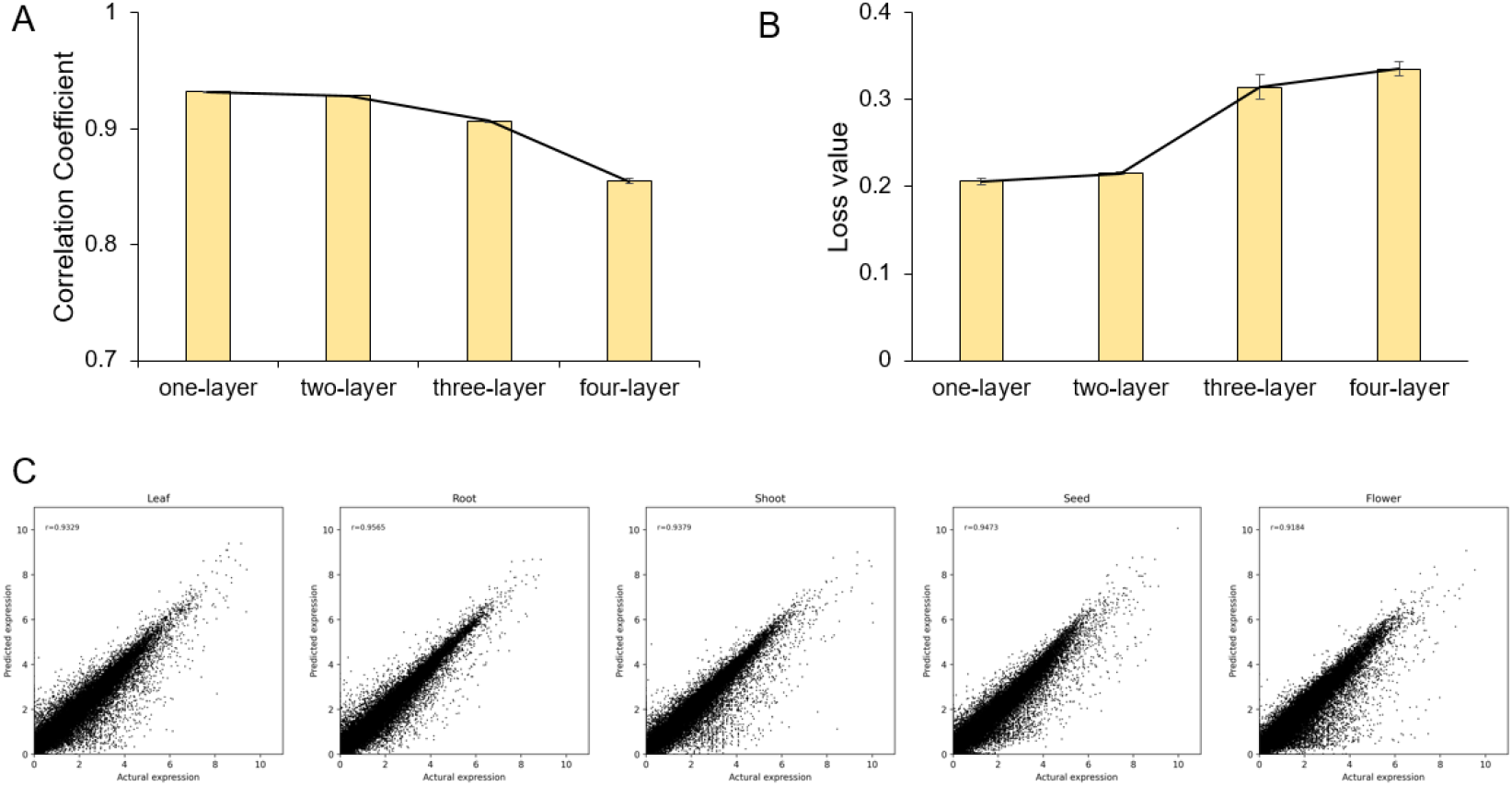
Gene expression prediction performance. (A) The predictive model’s performance with different layers. The correlation coefficient between precited expression levels and actual levels was calculated. (B) The predictive model’s loss value with different layers. (C) The model’s performances in soybean different tissues.

### Evaluate the predictive model’s performance

To test the generality of the prediction model, we used it to predict the expression of recently published independent data, which included mRNA profiles from four tissues: leaf, flower and pod ^23-25^. Using the expression levels of TF/TR genes, the predictor accurately reconstituted the expression of the other genes in high correlation with r = 0.9083 in leaf sample (n=6), r = 0.8889 in root sample (n=12), r=0.8820 in flower sample (n=3) and r=0.9022 (n=3) in pod samples (**Figure 3A-D, Figure S4**). The results indicated that even though gene expression levels varied considerably across samples in different experiments, our model is robust to predict the expression profiles.

**Figure 3.**
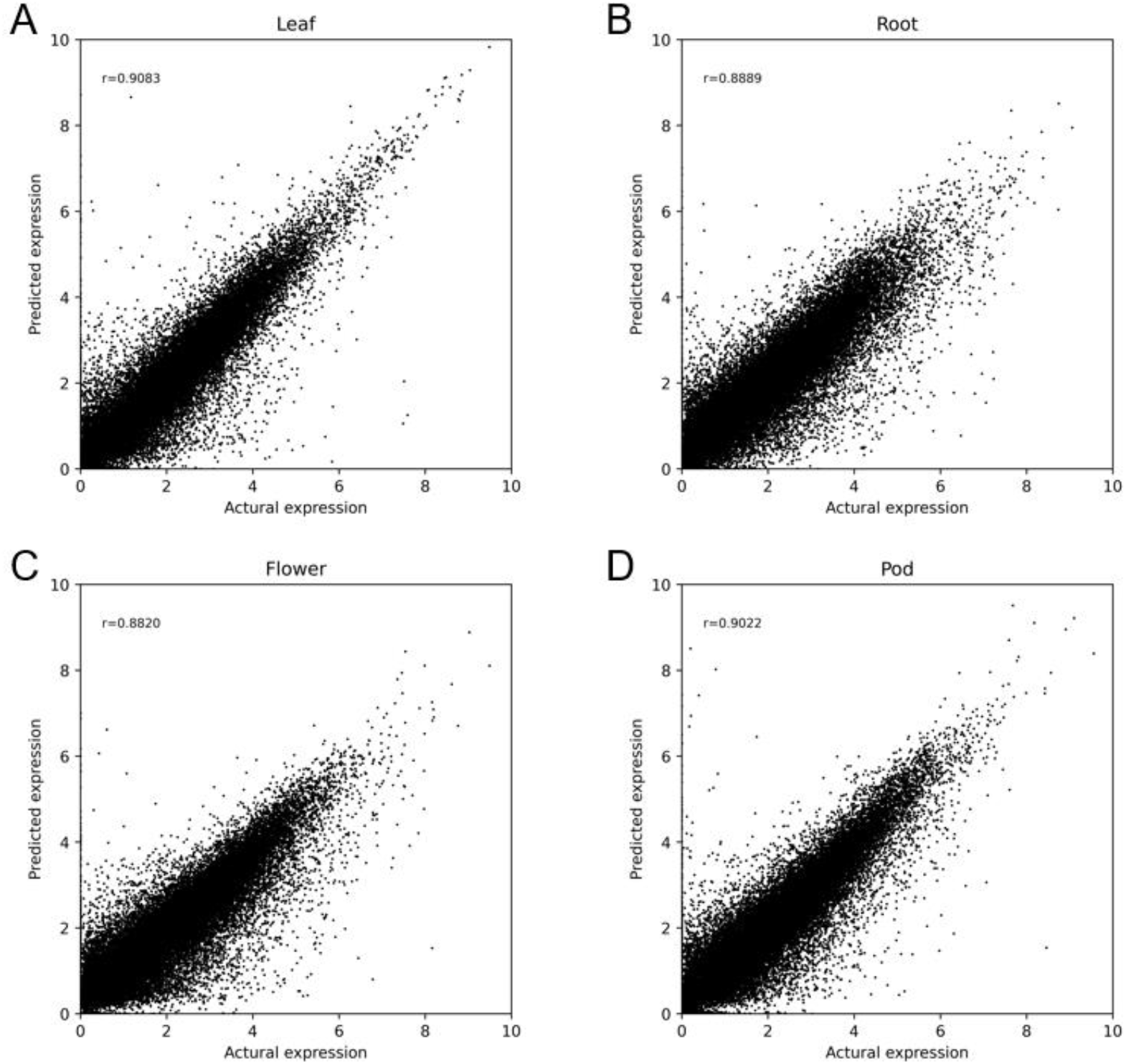
Prediction performance of the model on independent studies. The correlation coefficient of predicted expression and actual expression in leaf (A), root (B), flower (C) and pod (D) samples.

We continued to study the predictive ability on expression in untrained tissues: endosperm, epicotyl, petiole, and radicle ^15^. There was a strong correlation between the predictive expression generated by the model comparing to the actual expression (**Table 1**). While it is likely that nutrients such as starch is somewhat more in the endosperm thus affecting gene expression, these genes appear to be somewhat changed. This demonstrated that the model captured a broad representation of soybean tissues. In conclusion, our results suggested that tissue expression profiles can be effectively predicted based on the expression of TF/TR genes in the same tissues or developmental time points in soybean.

**Table 1.** Prediction performance of the model on unseen soybean tissues.

|  | **r** | **NRSME** |
| --- | --- | --- |
| endosperm | 0.8249±0.0162 | 0.3917±0.0298 |
| epicotyl | 0.8704±0.0214 | 0.3099±0.0382 |
| petiole | 0.9012±0.0083 | 0.2424±0.0107 |
| radicle | 0.8604±0.0134 | 0.3159±0.0337 |

### Identification candidate tissue-specific transcriptional regulators

We focused our analysis on the data of three tissues (leaf, shoot and root) which included sample number more 1000. We trained the XGBoost model to classify these tissues using TF/TR genes from leaf, root, and shoot samples, totaling 4017 samples. The model got a high accuracy (accuracy score: 0.99, precision: 0.98, recall: 0.99, F1 score: 0.98) using a 10-fold cross-validation. The loss curves show that the loss values decreased as the number of times increased, and the training and validation sets show the same trend (**Figure S5)**. After training, 744 candidate TF/TR genes were identified with feature importance more than 1, such as bHLH, ERF, MYB, NAC and WRKY TF genes. The top 20 TF/TR genes were showed in the **Figure S6**.

To validate the candidate TF/TR genes, we used the other remaining transcription factors to retrain our one-layer predictive model. The results showed that the accuracy of the model increased to 0.9438% and the loss value decreased to 0.1904. And the test accuracy of the model in the root sample increased to 91.03%. It supposed that these candidate TF/TRs genes were the key genes that played important roles in the development of these tissues.

### Degree centrality analysis revealed hub modules in tissues

The co-expression networks were constructed on the correlations among genes according to the trends of gene expression in all soybean tissues. Genes with high correlation coefficients that indicated a high degree of interconnections between the genes were defined as modules with each module depicted by a branch of different color and each gene depicted by a leaf. A total of 316 modules were identified. After hierarchically clustering the genes based on TOM dissimilarity and merging the modules with at least 80% similarity, we generated a GCN containing 31 merged modules (**Figure 4A**). The numerical representation of the correlation coefficient between modules and different tissue types can be used to identify the tissue-specific modules (**Figure 4B**). Notably, five co-expression modules are highly expressed in the leaf, including MEmistyrose, MEsienna2, MEhotpink2, MEburlywood2 and MEthistle2 modules. In the root, 12 co-expression modules were highly expressed, including MEcoral2, MEchocolate1, MEgoldenrod4, MEbisque, MEdeeppink, MElightpink3, MEhotpink1, MEburlywood, MEgrey, MEturquoise2, MEhoneydew and MEdarkorange modules. There were no strongly correlated co-expression modules in seedling, flower, hypocotyl, and cotyledon. This may be due to the small sample size of these tissues.

**Figure 4.**
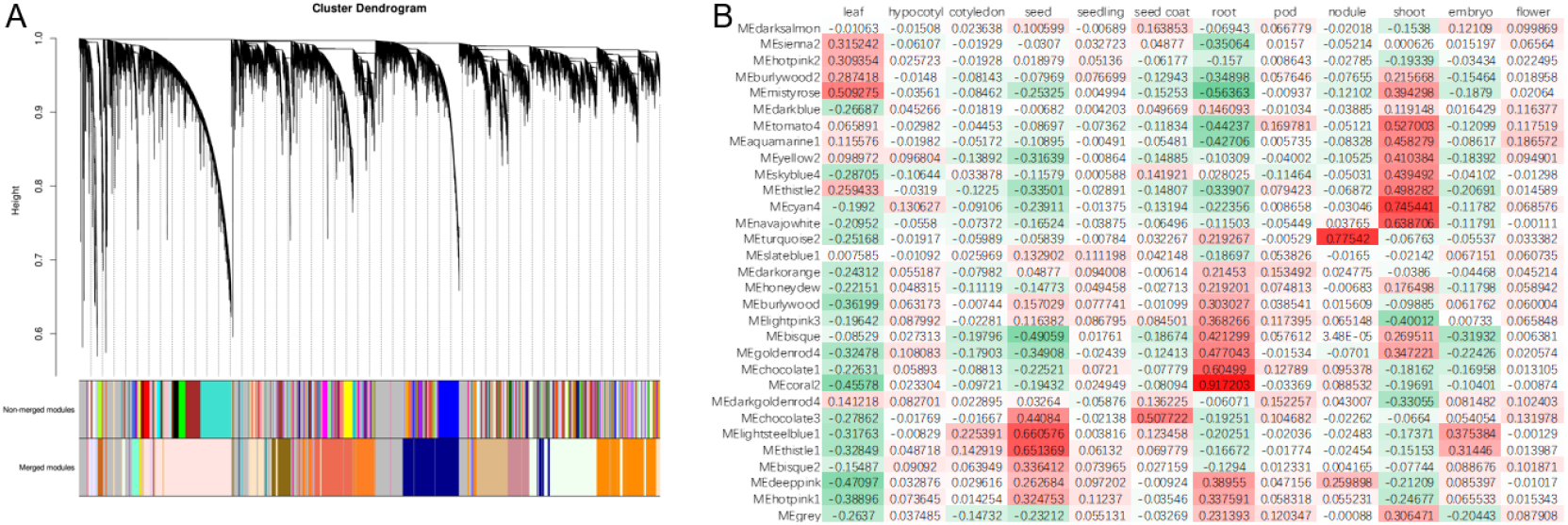
Dendrogram and module colors before and after merging similar modules. (A) Hierarchical cluster tree showing co-expression modules identified by WGCNA analysis. Each leaf in the tree is one gene. Modules were merged based on similarity of their eigengenes. The major tree branches constitute 31 modules labeled by different colors. (B) Module-tissue association. Each row corresponds to a module. Each column corresponds to a specific tissue. The color of each cell at the row-column intersection indicates the correlation coefficient between the module and the tissue type. A high degree of correlation between a specific module and the tissue type is indicated by red or green.

### Identification TF-regulated modules

We next combined the gene co-expression modules and these candidate TF/TR genes. In the leaf, 5836 genes were highly specifically accumulated in the MEmistyrose module, which indicated that this group of genes might be responsible for leaf development. GO analysis showed that they mainly enriched in response to photosynthesis and response to chemical stimulus, which supposed the leaf responds the complicated light conditions (**Figure 5A**). A total of 2842 genes involved in MEcoral2 module were highly specifically accumulated in the root, which indicated that this group of genes might be involved in the developmental process in root. GO analysis showed that they mainly enriched in response to stimulus and metabolic progress, which supposed the different responses of roots to complicated soil environments (**Figure 5B**).

**Figure 5.**
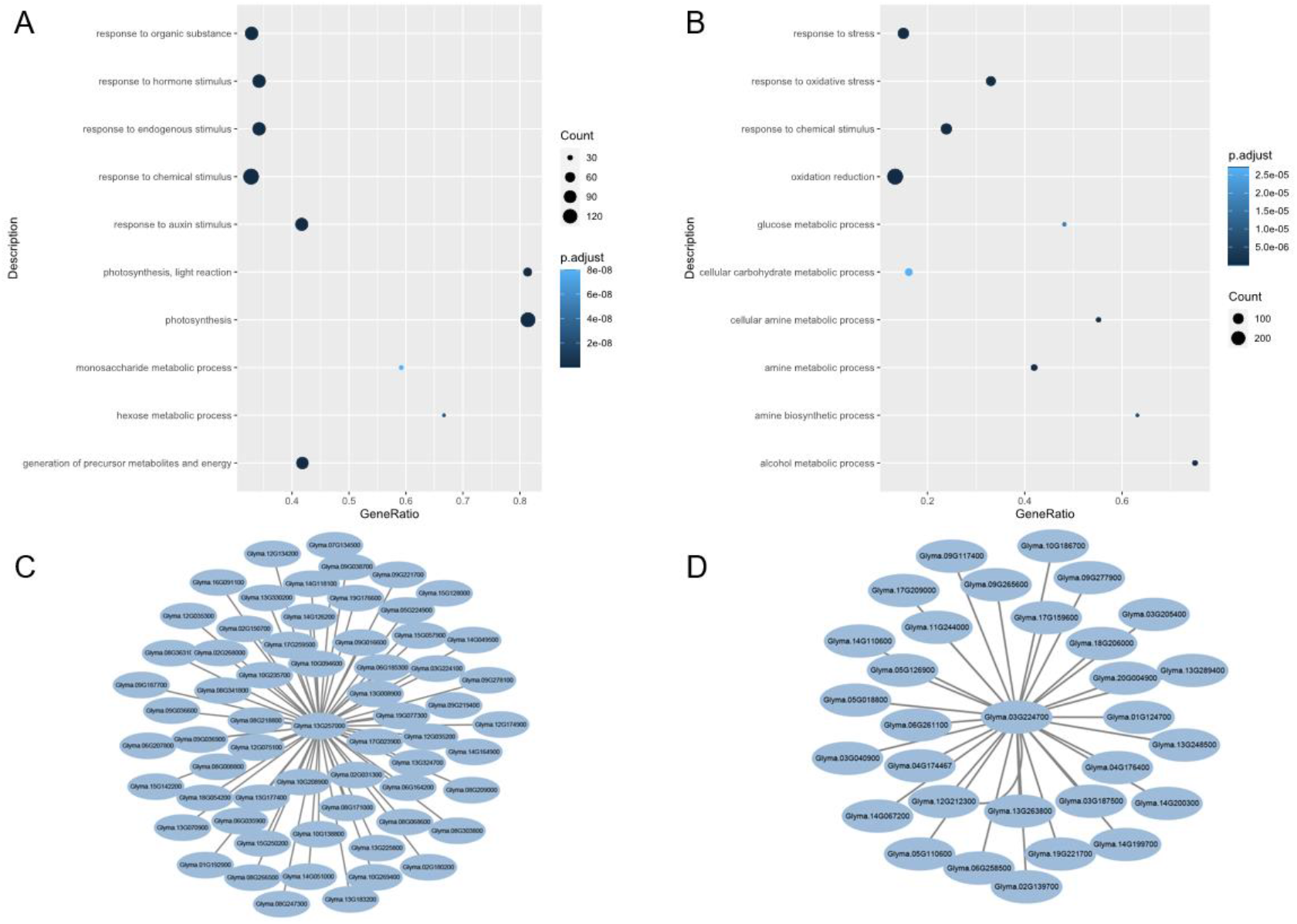
Modules with enriched biological (GO) processes. (A) GO enrichment analyses of 5836 genes in the MEturquoise module. (B) GO enrichment analyses of 2842 genes in the MEcoral2 module. The top 10 terms for biological progress in the GO category were showed. (C) A MADS TF gene (Glyma.13G257000) regulatory network in co-expression modules. (D) AMADS TF gene (Glyma.03G224700) regulatory network in co-expression modules.

The regulatory relationship between the TF/TR genes and target genes in the modules were inferred by GRNBoost2 algorithm and top 1% connectivity were elected by the importance (**Table S1, Table S2**). For example, a MADS-box (*Glyma*.*13G257000*) and a WRKY (*Glyma*.*03G224700*) gene regulatory networks were showed in **Figure 5C, 5D**. *Glyma*.*13G257000* is a homolog of *AtAGL22*, which was a central transcription factor for balancing developmental and stress tolerance processes in *Arabidops* ^35^. It interacted with E1 (Glyma.06G207800), which was an important legume-specific transcription factor that was involved in the regulation of leaf morphology in collaboration with other TFs ^36^. GmWRKY27 (Glyma.03G224700) was found to be involved in stress tolerance in soybean root ^37^. This suggested that the transcriptional regulatory module centered on TFs/TRs may offer great convenience and direction over traditional RNA assay analyses in providing regulation of developmental traits.

## Discussion

Here, we proposed a combination of TF/TR gene expression data to characterize the universality of genes within soybean. The classification model was then used to identify tissue-specific candidate TF/TR genes, this ranking method could help to reveal candidate genes and signatures of selection that may explain differences in the developmental processes of different tissues. Although the phenotypic consequences of differences in the function of a gene in different tissues remain to be determined, we found that by identifying a number of specific gene regulatory modules that vary across tissues, this suggests that these changes may have some phenotypic effects. Overall, these findings provide new insights and new candidate genes for the direction of TF/TR regulation of gene expression in soybean research. Future work would be required to determine how these modules occurred in the soybean development.

While we have shown that our method can aid in the analysis of soybean expression data, its applicability to different tissues underlies its potentially broad utility. Importantly, the strong correlation between the predict expression in unseen data in root, flower and pod suggests that TF/TR genes measured in different tissue types can unearth the homeostasis of target genes (**Table 1**). For those large library samples, predictive models have been applied in Arabidopsis and human with excellent results ^16, 17^. However, there is a risk of overfitting when the training samples are less than 20,000 (**Table S3**). Our results show that even with a relatively small sample size in non-model species, soybean, our method can still capture meaningful gene expression information. Our approach has the potential to aid in identifying patterns of gene expression regulation in different tissues small samples.

We also investigated the tissue-specific gene modules through all the transcriptomes of different tissues of soybean. Highly expressed modules were found in leaf, seed, seed coat, root nodule and shoot (**Figure 4B)**. It showed that gene co-expression modules played important roles in the process of soybean growth and development, which is consistent with the results of the previous study ^14^. The plants developed a pattern of sophisticated regulation of genes present in different tissues. These modules were found to contain many TFs, suggesting that these modules may correspond to systems that rely on more complex transcriptional regulatory structures.

By exploring key TF/TR gene sets in tissue classification, we combined these TF/TR gene with co-expression modules. The TF/TR genes and their target genes were identified. In the soybean, E1 was found to regulate flowering time in soybean and has been domesticated by humans over a long period of time ^38^. And it could also be found to be involved in regulating the leaf traits with different TF genes ^36^. In the leaf module, the interacting could also be found. In root, GmWRKY27 gene involved in salt and drought tolerance in soybean root hair development ^37^. GmMYB29A2 (Glyma.02G005600) gene regulated the soybean root hair resistance to pathogen ^39^. GmWRKY6 (Glyma.15G110300) was involved in the progress during the low phosphorus, and GmNAC109 (Glyma.14G152700) gene control the lateral root formation and abiotic stress ^40, 41^. Shoot has important roles in nutrient transportation and supporting leaves in plants. In the shoot modules, *GmAP1a* (*Glyma*.*16G091300*) is involved in a biological process that regulates the length of soybean shoot ^42^. GmIAA27 (Glyma.09G193000) encoded an important AUX/IAA protein and overexpression of the mutant type of this gene in *Arabidopsis* resulted in dwarf phenotype ^43^. These genes could be found in these corresponding co-expression modules. Thus, by using these transcriptional regulatory modules centered on TFs/TRs in soybean will be an exciting area for further research.

## Conclusion

We outlined a strategy that uses TF/TR gene expression data from transcriptome studies to prioritize genes that are more likely to conserved genes in the developmental homeostasis in soybean and applied this method with classified model to identify candidate TF/TR genes related to soybean tissue development. Our findings provide opportunities for targeted follow-up experiments and increase our understanding of how TF/TR genes have shaped soybean tissue development. Overall, we anticipate that our method will become a useful tool for identifying functionally significant gene expression changes in many tissues across diverse non-model species and will contribute to our understanding of how gene expression drives diversification of traits.

## Acknowledgements

This research was supported by Guangzhou University Graduate Student Innovation Ability Cultivation Program (2022GDJC-D14).

